# Shrimp Hemocyanin elicits a potent humoral response in mammals and is favorable to hapten conjugation

**DOI:** 10.1101/2024.01.11.575311

**Authors:** Huiwen Sun, Moris Wei, Amber Guo, Ci Zhang, Yuefeng Wang, Renhui Huang, Xiaoxiao Li, Jeffrey Zhan, Jonny Wu, Bruce Jiang

## Abstract

Conjugation to a carrier protein is essential to give rise to the antigenicity of hapten, which is a small molecule and cannot induce an immune response by itself alone. Three carrier proteins e.g. KLH (Keyhole Limpet Hemocyanin), BSA (Bovine Serum Albumin), and OVA (Ovalbumin) were used mostly. KLH is advantageous to the others, majorly owing to its strong immunogenicity and limited usage in other biological assays. However, the solubility of KLH is not as well as the others, especially after hapten conjugation. Besides, the high market price of KLH results in high costs in vaccine and antibody development. Here, we extracted the shrimp hemocyanin (SHC) from *Penaeus vannamei (P. vannamei)* with a production yield of > 1 g proteins (98 % pure) per 1 kg shrimp. Compared to KLH, the peptide-SHC conjugates exhibit higher solubility after hapten conjugation. Furthermore, compared with KLH, SHC induces comparable antibody production efficiency in mammals, with or without conjugation. Finally, rabbit polyclonal antibodies or mouse monoclonal antibodies were generated by immunizing SHC-peptide conjugates, and their applications in western blot, immunofluorescence and immunohistochemistry were confirmed. Therefore, we demonstrated that SHC may be used as a substitute for KLH in future antibody and vaccine development.

## Introduction

Hapten, a low molecular weight substance without immunogenicity, could stimulate immune response and antibody production when coupled to a carrier protein (1-3). The mechanism may be explained by that hapten-carrier conjugate is able to cross-link B-cell receptors and activate T cells through the MHC class II presentation (4, 5). For the generation of antibodies against hapten or the hapten vaccine, three carrier proteins were used for covalent conjugation, e.g. KLH (Keyhole Limpet Hemocyanin), BSA (Bovine Serum Albumin) and OVA (Ovalbumin) (6). BSA and KLH could induce a strong immune response, while OVA is less effective and its solubility is not as high as the others (7). The prevalent usage of BSA in many other biological assays has limited its use for immunization. For instance, BSA is usually used as a blocking reagent in Elisa, where anti-BSA antibody would interfere with the downstream detection. KLH is regarded as the priority for preparing hapten-carrier conjugates for immunization. However, its solubility and cost are not as well as BSA, especially after peptide conjugation. KLH is purified from the hemolymph of the inedible marine mollusc, Keyhole limpet (*Megathura crenulata)* native to the Pacific coastal waters of California and Mexico (8). Aquaculture of Keyhole limpet is the main way to provide the animal resources for KLH extraction on a large scale, which results in the high economical cost for antibody or vaccine development. Therefore, an appropriate substitute with better biochemical characteristics, and lower economical cost, is still needed for preparing hapten-carrier conjugates.

Hemocyanins are respiratory proteins occurring freely dissolved in the hemolymph of many arthropods and molluscs (9-11). Similar to molluscs, arthropods are evolutionally distant from mammals (12), suggesting that hemocyanin from arthropods may induce a strong immune response in mammals. The hemocyanin from the white-leg shrimp (*Penaeus vannamei (P. vannamei)*) (13)(14), which belongs to the phylum Arthropoda, was considered after several logical thoughts. Firstly, *P. vannamei* is a traditional marine food and its aquaculture is prevalent worldwide. According to the Food and Agriculture Organization (FAO) of the United Nations, aquaculture of *P. vannamei* contributed to 80% of the whole shrimp production, which reached 4, 000, 000 metric tonnes in 2014 (15). This data means the obtainment of *P. vannamei* is easy and economical. Secondly, the molecular weight of the arthropod hemocyanin subunit is 72kDa, which is much lower than KLH (350 or 400kDa in molecular weight) (9). In general, lower weight means a higher solubility. In this study, we extracted hemocyanin from *P. vannamei* and determined its potential to serve as a peptide-conjugated carrier protein for antibody development.

## Materials and Methods

### Reagents

Sulfosuccinimidyl-4-[N-maleimidomethyl] cyclohexane-1-carboxylate) (sulfo-SMCC) (*Cat.* M123456), 1-ethyl-3-(3-dimethylaminopropyl)carbodiimide hydrochloride (EDC) (*Cat.* A638729) were purchased from Aladdin (Shanghai). KLH (Cat. 77600) was purchased from ThermoFisher. Peptide synthesis was conducted by Genscript (Nanjing) in the form of dried powder. The sequence of Peptides is shown in the supplementary table.

### Preparation of Shrimp Serum

Healthy *Penaeus vannamei* with a weight between 15–20 g from Shanghai Didong Corporation were obtained and reared in 25-L seawater tanks at 25°C. Air was continuously supplied using an electric pump. Hemolymph was taken directly from the pericardial sinus using a sterile tube and then allowed to clot overnight at 4°C as described in previous study (16). The serum was separated after centrifuging at 5,000 *g* for 10 min and kept at −20°C until use. All animal experiments were conducted according to the recommendations from the Animal Ethics Procedures and Guidelines of the People"s Republic of China.

### Chromatographies

SHC purification was performed by gel-filtration chromatography and anion-exchange chromatography as previously described (16). Briefly, 2 ml of *L. vannamei* serum was loaded onto a Sephadex G-100 column, and then, the column was washed with Tris–HCl buffer (0.05 M, pH 8.0) at a flow rate of 1 ml/min until the absorbance at 280 nm reached baseline. Then, the above proteins were loaded onto a DEAE-cellulose column, equilibrated with 0.01 M pH 7.5 phosphate-buffered saline (PBS) buffer. Elution was performed with 0.5 M NaCl at a flow rate of 2 ml/min. proteins concentration was determined by a nano-drop at OD280 and the purity was detected by commassie-blue (Epizyme) staining after SDS-PAGE separation.

### Peptide conjugation

Carrier protein dissolved in phosphate-buffered saline (PBS) at the concentration of 10 mg/mL was mixed with 1mg/mL sulfo-SMCC in PBS at room temperature for 1 h. Then, the mixture was loaded on a Sephadex G25 column for desalting using 0.1M sodium phosphate buffer, pH 7.0. Activated protein was collected according to the optical density (OD) of 280 nm and 260 nm (ratio of 280/260 > 1 because SMCC has a high absorbing density of OD260 compared to protein). Next, the peptide dissolved in PBS was mixed with the activated carrier at the final concentration of 1 mg/mL at room temperature for 2 hours. For EDC conjugation to prepare peptide-conjugates in Elisa screening, 1 mg/mL EDC was mixed with 10 mg/mL BSA and 1mg/mL peptide, which were dissolved in MES pH3.5 buffer at room temperature for 30 min.

### Immunization

Rabbits were immunized by administering biweekly injections of the antigen with a total amount of 1mg in adjuvant. The injection sites on the rabbit are shaved and disinfected before immunization. Complete Freund’s adjuvant is only used with the first immunization, and incomplete Freund’s adjuvant is used with the secondary immunization. Subsequent immunizations are performed in phosphate-buffered saline (PBS) without Incomplete Freund’s adjuvant for boosting. Serum samples of 40 mL were collected after the termination of immunization, normally 30 days after the first immunization. Similarly, the mouse BALB/c were immunized with 10 ug antigen biweekly and the blood sample was collected at a 3-day interval. This experiment was conducted with the help of Qingdao Kangda Biotech.

### Hybridoma fusion and mono-antibody screening

To isolate B-lymphocyte producing certain antibodies, BALB/c female mice are immunized through repeated injection of a specific antigen. The intended result is the formation of hybridoma cells formed by fusion of B-cell and myeloma cells which is done by using Polyethylene glycol (PEG) as described previously (17). The hybridoma cells were selected under HAT medium, and each supernatant for antibodies by Elisa. Screening is done to select hybridoma cells which are the desired cells for monoclonal antibody production. Subsequently, the selected hybridoma cells are cultured in a suitable culture medium for antibody production in vitro.

### Cell culture

Cells were cultured in DMEM medium supplemented with 2 mM L-glutamine, nonessential amino acids, 100 U penicillin per mL, 100 μg streptomycin per mL and 10% FBS (complete DMEM). Cells were passaged after the confluence reached 90%. All of the cell lines were a kindly gift from Shanghai Fuheng Biotech.

### Western blot analysis

Western blot analysis was performed as previously described (9). Briefly, samples were collected in a loading buffer (Epizyme) after one wash with PBS, and total proteins were separated by SDS polyacrylamide gel electrophoresis and transferred onto an NC membrane. The membrane was blocked using 5% dried milk and incubated with the indicated primary antibody and secondary antibody. Bands were visualized using Omni ECL reagent (EpiZyme) under an illuminating imager XF101 (Epizyme), and the grey intensity was acquired by using Fiji (NCBI).

### Immunohistochemistry (IHC)

Tissue was fixed with formaldehyde for 3 hours at room temperature, then dehydrated at a sequential concentration of ethanol and embedded in paraffin. The samples were then sectioned at a thickness of 5 to 20 μm and paraffin was removed by xylene and hydrated at a sequential concentration of ethanol. Antigen retrieval was by heat mediation in Tris pH 9. Samples were blocked with 1% BSA and incubated with primary antibody for 16 hours at 4℃. An HRP-conjugated goat anti-rabbit IgG polyclonal was used as the secondary antibody. DAB was added to develop the signal for 1-5 mins until the brown color in the sample. The remaining DAB was removed by water several times, the nuclei were further stained by Hematoxylin. The sample was mounted in a resin for photography.

### Immunofluorescence

Cells were fixed with 4% paraformaldehyde for 5 mins and penetrated with 0.05% TrionX-100 for 2 mins at room temperature. Then, 10% FBS in PBS were used for blocking, and the primary antibody was treated for 1 h, followed by the FITC-conjugated goat anti-mouse IgG at a dilution of 1: 5000 for 1h. Nuclei were counterstained with DAPI for 1 min. The image was captured by KEYENCE BZ-X800LE through DAPI and GFP channels and data were generated by deconvolution process.

### Electron-microscopy (EM)

EM was performed as describe previously (18). In brief, 3ul protein (10 ug/uL) dissolved in 1× PBS was dropped on a piece of para-film, and a carbon-coated nickel grid was placed on the drop for 30–60 minutes. The sample was treated with 2% uranyl acetate for 30 seconds, washed three times with water. The grid was air-dried and disinfected under UV (ultra-violet) before examination by FEI F20 TEM at 120 kV.

### Statistical analysis

The continuous variables in different subgroups were compared using an unpaired t-test and a one-way analysis of variance. All the tests were two-sided, and p < 0.05 was considered statistically significant. The statistical analyses were performed using GraphPad.

## Results

### High pure SHC was acquired from *P. vannamei* by two-step chromatography

Adult *P. vannamei* obtained from the market was applied to hemolymph extraction. Then, serum containing SHC was collected after the centrifugation of clotted hemolymph and SHC was further purified by chromatography (Figure 1A). One shrimp is about 10g in weight on average, which could provide 0.5 mL of whole hemolymph after extraction through the pericardial sinus region (Figure 1B). The hemolymph and serum showed a clear black-blue color, indicating that SHC is rich in the liquid (Figure 1B). In addition, SHC could be visualized by coomassie-blue staining after SDS-PAGE separation, where a band with a molecular weight close to 70 kDa was observed (supplementary figure 1). Next, we conducted the purification via size exclusion and ion exchange chromatography. After the two-step purification, SHC with 98% purity is acquired (figure 1C, 1D). On the other hand, the solubility of SHC is about 200 mg/mL, which is comparable to its counterpart KLH (supplementary figure 1B). Furthermore, size-exclusion chromatography analysis showed that 80% of SHC were between 158 kDa to 440 kDa in weight, while the remaining between 44 kDa to 75 kDa (figure 1E), suggesting that SHC preferred to form polymers in physical environment. Consistent with this, we observed that most proteins were nearly 20 nm in diameter and a small amount of them were less than 10 nm under transmission electron microscope (figure 1F), which corresponded to the diameter of the monomer or tetramer simulated by Alpha-fold2 based on the primary amino-acids sequence (figure 1G). Overall, we could postulated that SHC is able to polymerize to form the tetramer *in-vitro*. Altogether, the production efficiency of high pure SHC protein is more than 1 g per 1 kg shrimps, highlighting the possibility of large-scale SHC production for commercial usage.

**Figure 1.**
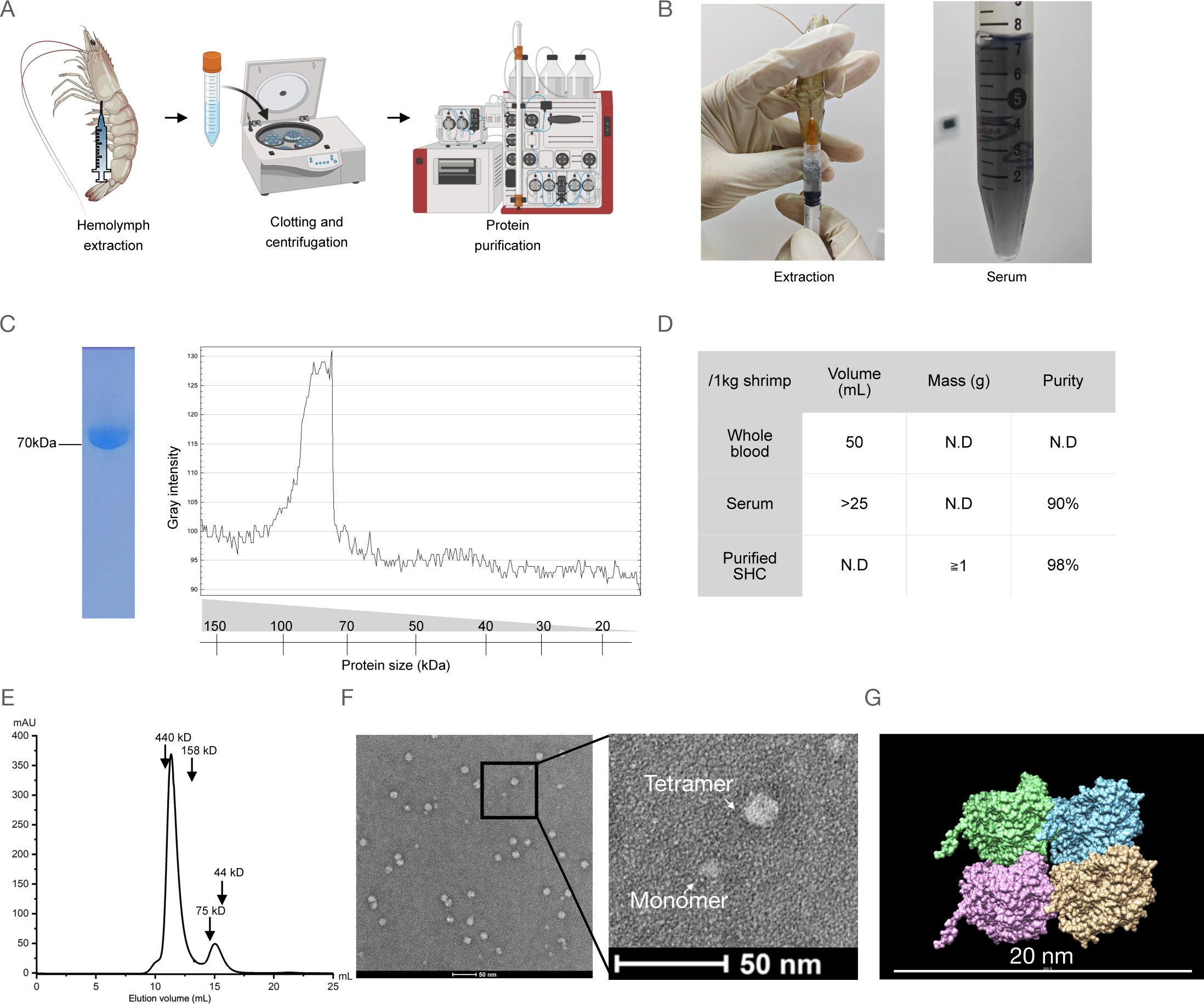
SHC extraction and purification from *P. vannamei.* (A) Schematic representation of the SHC purification. (B) Photos of hemolymph and serum extracted from shrimps. A blue-black color could be observed. (C) SDS-PAGE separation of purified SHC. Proteins were detected by Commassie-blue staining. Fiji software was used to acquire the grey intensity in the gel. (D) A calculation of the SHC yield and purity at the indicated steps. N.D. represents not determined. (E) Size exclusion chromatography of purified SHC. SHC was dissolved in PBS and loaded into Superdex-200 column. Size was determined by running of standard proteins representing different molecular size. (F) Electron-microscopy analysis of purified SHC. (G) Alpha-fold2 prediction of SHC protein monomer. Four colors represented the four units of SHC.

### SHC is superior to KLH in solubility when conjugated to peptides

Among 672 amino acids (Supplementary Figure 2A) in SHC, there are 36 Lys (K), 57 Asp (D), 47 Glu (E) and 2 Cys (C). Thus, besides the terminal -NH3 and -COOH, one SHC contains 36 -NH3, 104 -COOH and 2 -SH. Alpha-fold2 structural simulation of SHC indicated that there are adequate Lys, Asp and Glu exposed on the surface (Figure 2A), which may be accessible for hapten linkage. We applied a conjugation assay to test the conjugation efficiency of SHC, where FITC-labelled peptide and SMCC linker were explored (Figure 2B). In the neutral pH environment, SMCC activates -NH3 in the donor, and the SMCC-activated SHC was desalted by a G25 column (Supplementary Figure 2B). The activated amine then selectively attacks -SH in the recipient and forms a covalent bond (Figure 2B). Of note, in the reaction mix, 1 mg/mL SMCC-activated carriers (BSA, 15.3 μM; SHC, 13.8 μM; KLH, 2.8 μM) were mixed with 1 mg/mL FITC-peptide (1.68kDa, 618 μM) in their final concentration. After the conjugation, 2.5 uL conjugates were loaded into SDS-PAGE gel. Signals were detected by the fluorescence via a 488 nm filter (Figure 2C). BSA, KLH and SHC formed the covalent bond to FITC-labelled peptide with different binding efficiency. About 35.46% of peptides conjugated to the BSA were soluble (Figure 2C, 2D). About 33.09% of peptides linked to SHC were soluble and 6.77% were insoluble. In contrast, insoluble KLH-peptide conjugates dominate, as only 1.51% were soluble. Including soluble and insoluble conjugates, the conjugation efficiency of BSA, SHC and KLH was calculated as 35.46%, 39.86%, and 23.54%, respectively, suggesting that one molecule of BSA, SHC and KLH is able to link approximately 14, 17 and 59 peptide molecules at average, respectively. In principle, one SHC may link 36 peptide molecules. This difference between theoretical prediction and experimental results may result from the steric hindrance or the protein cross-linking. Nevertheless, these data suggested that SHC is suitable for hapten conjugation and its solubility is superior to KLH after peptide conjugation.

**Figure 2.**
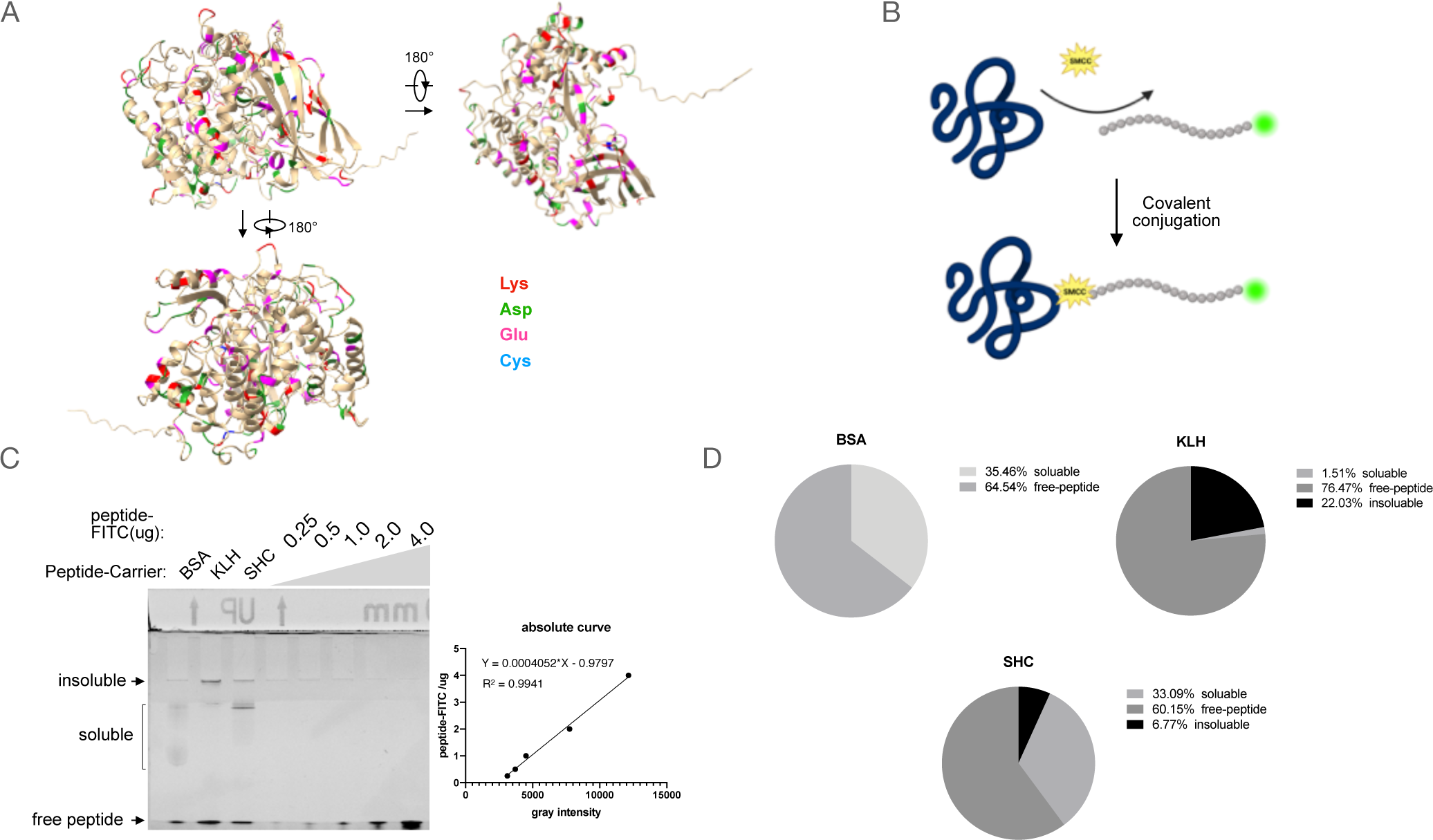
Generation of SHC-peptide conjugates. (A) 3D structure of SHC proteins predicted by Alpha-fold2. (B) Schematic representation of SMCC-mediated covalent linkage of peptide to carrier proteins. (C) SDS-PAGE gel separation of peptide-carrier conjugates. A serial dilution of FITC-peptide was used to make the standard curve for the absolute determination of conjugates. Of note, a large complex that stayed in the sample well was defined as the insoluble conjugates, which were turbid in reaction liquid. (D) Statistics of conjugation efficiency and conjugates solubility.

### The immunogenicity of SHC is potent in mammals compared with KLH

To determine the immunogenicity of SHC in mammals, equal amounts of KLH and SHC were given to BALB/c mice. Blood sample were harvested every three days and the titer of anti-KLH or anti-SHC were determined by Elisa assay. Humoral antibody response could be detected fourteen days post the first immunization, while strong antibody production against KLH or SHC was seen thirty days post the first immunization (Figure 3A). No significant difference in antibody titers could be observed between KLH and SHC immunized groups. Furthermore, rabbits were immunized by three carriers (KLH, SHC and poly-lysine) conjugated to the peptide designed for the PDI gene. Poly-lysine was set as a negative control as it generally does not induce a strong immune response in mammals. The rabbit serum was collected and applied to WB analysis, where cell-line samples were used as the native antigen (Figure 3B). Note that equal cells were loaded and no difference in the mass of whole cellular proteins was observable as confirmed by commassie-blue staining (data not shown). Consistency with the transcriptional level of PDI mRNA (supplementary figure 3), a similar protein expression pattern of PDI was detected in HEK293T, HeLa and HepG2 by the serum from rabbits immunized with KLH or SHC conjugates. Together, we demonstrated that SHC owns the potent immunogenicity compared with KLH.

**Figure 3.**
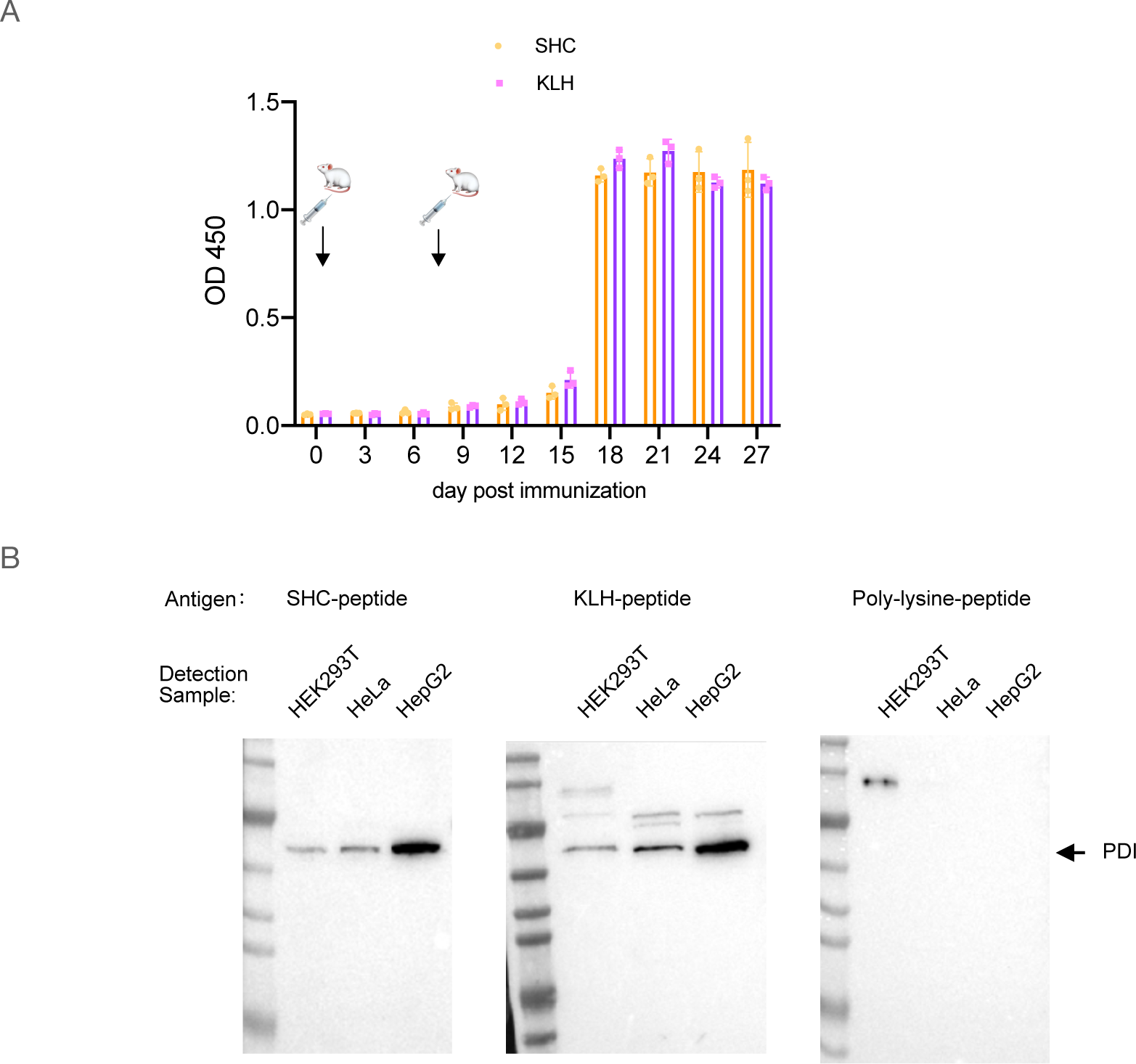
SHC owns the comparable immunogenicity to KLH. (A) Titer determination of humoral response after mice immunized by SHC and KLH. In each group, 3 female mice were used for immunization at the indicated time in figure. Elisa was used to detect the level of anti-SHC or anti-KLH antibodies. (B) SHC-peptide stimulated the production of rabbit antibodies against the PDI gene. Equal cell numbers were loaded for WB detection.

### Confirmation of SHC extensiveness for rabbit polyclonal antibody development

To extend the use of SHC for hapten immunization, SHC-peptide conjugates were prepared for generating poly-antibody against the following several genes: α-tubulin, α- actin, β-actin, Desmin, NF-κB, Cytokeratin 10, NRF2, IGF2R, most of which are cellular host genes or the common targets in biological research. WB and IHC were conducted to test the rabbit serum harvested thirty days after the first immunization. The rabbit sera against α-tubulin, β-actin, Desmin, and Cytokeratin-10 were applicable in both WB and IHC detection, while sera against others were applicable for at least one of the two assays (Figure 4A, 4B). WB data showed that the bands of cellular host genes were clear and antibody specificity was high. Although detection results of Desmin, NF-kB and Cytokertin-10 showed some un-specific bands, the specific bands are present in the correct molecular weight and the expression pattern was in consistence with their RNA transcription level (Supplementary Figure 4). On the other side, the positive signal in IHC analysis showed the correct morphological distribution of the corresponding antigen in tested organs (Figure 4B). Of note, no signal in control without the addition of primary serum could be detected (data not shown).

**Figure 4.**
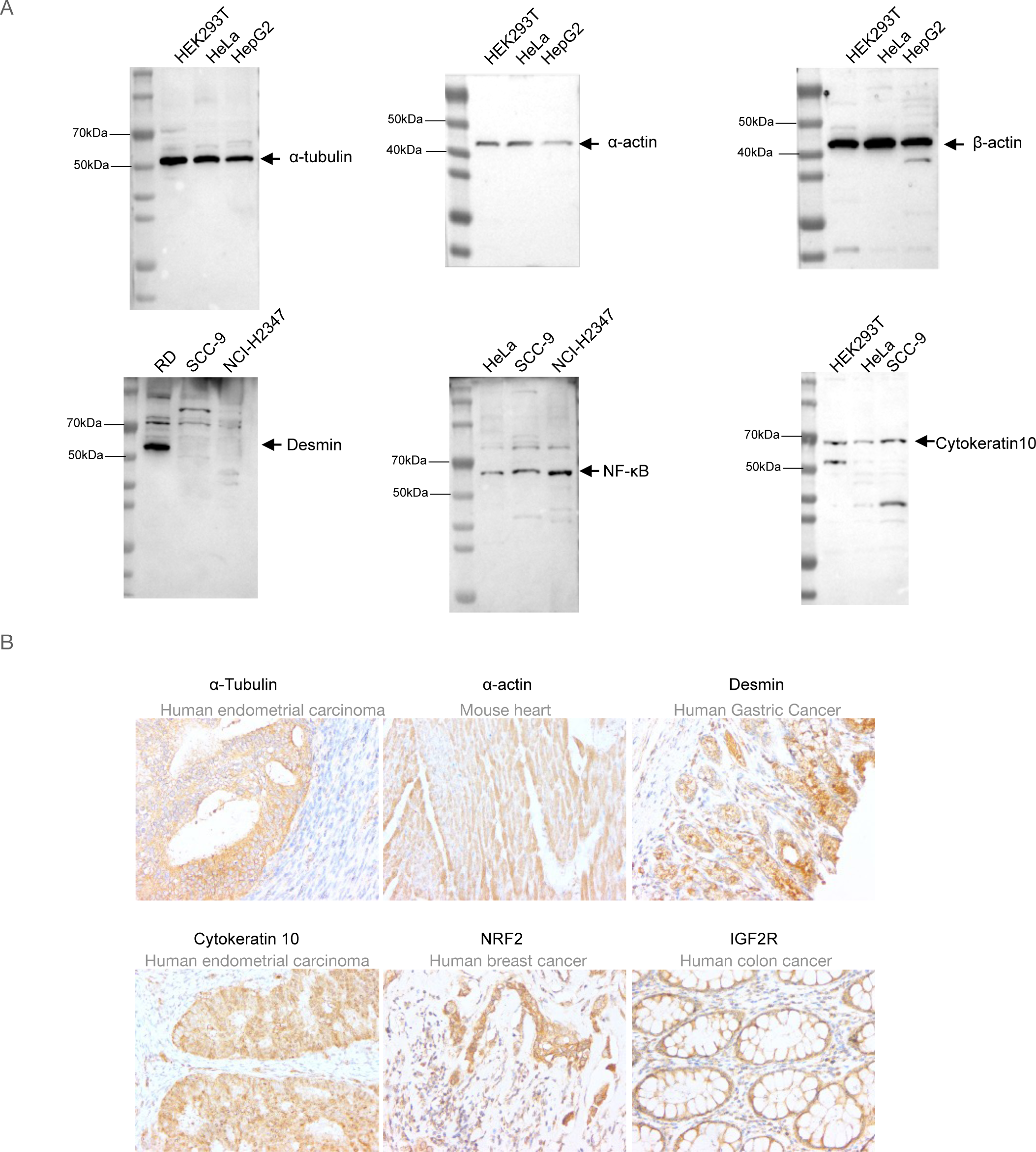
SHC could be used for rabbit polyclonal antibody development. Peptides targeting the indicated proteins were designed and conjugated to SHC. Rabbit serum was harvested 30 days after the first immunization. (A) Confirmation of the rabbit serum application in WB detection. Of note, equal cell numbers were loaded. (B) Confirmation of the rabbit serum application in IHC detection.

### Confirmation of SHC for application in mouse monoclonal antibody development

Next, we asked whether SHC could be applied in monoclonal antibody development. After immunization of peptide-SHC conjugates for anti-β-actin generation in BALB/c mouse, spleen were sectioned and splenocyte was isolated by ficol density centrifugation. PEG1450 were used for the fusion of Sp2/0 and splenocyte. Several positive clones were finally screened by Elias assays. One clone anti-β-actin (4B6) was purified from the hybridoma supernatant by protein G purification with the affinity of Kd = 10^-9^ M (figure 5 A, 5B). This 4B6 clone was further confirmed in WB application, where a cellular actin band close to 40 kDa was visualized (figure 5C). Filament-like morphology of actin in cytoplasm was clearly observed by immunofluorescence (figure 5D). Together, these data demonstrated the applicability of SHC in monoclonal antibody development.

**Figure 5.**
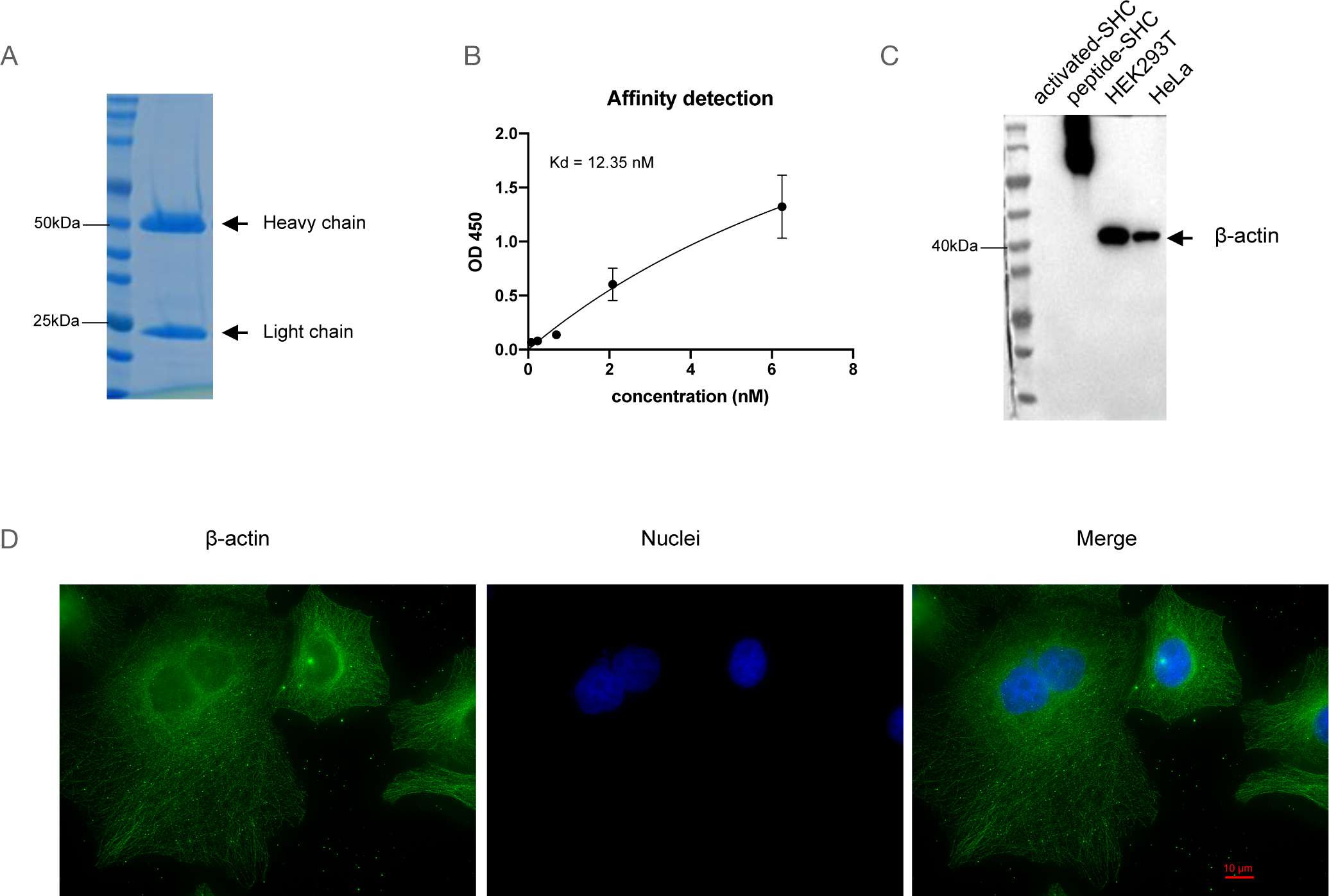
SHC could be used for mouse monoclonal antibody development. (A) Protein G column purification of anti-β-actin from hybridoma supernatant. The purified antibody was detected by SDS-PAGE separation. (B) Determination of kinetic dissociation (Kd) of peptide and anti-β-actin. Peptide-BSA conjugates were coated in 96-well, then a serial dilution of purified anti-β-actin were incubated as the primary antibody. All the other steps were performed as same as the Elisa assay. The Non-linear fit by one-site specific binding in GraphPad Prism 8 was applied to calculate the Kd value. R^2^ = 0.9345. (C) Confirmation of the application of anti-β-actin in WB detection. Peptide-SHC conjugates were set as a control. Equal cell numbers were loaded. (D) Confirmation of the anti-β-actin application in IF detection. β-actin was visualized by the green while nuclei were visualized by blue. HeLa was used for this detection.

## Discussion

Carrier proteins play a vital role in the generation of hapten-related antibodies and vaccines because hapten is not able to be processed by antigen-presenting cells with itself alone. Nowadays, KLH is the most commonly utilized carrier due to its strong immunogenicity and limited application in biochemical assays. However, obtainment of KLH mainly relies on the aquaculture of the sea animal keyhole limpet, which is generally inedible worldwide. Moreover, the solubility of KLH is low after conjugation, resulting the difficulty in the downstream purification. Here, we determined that SHC possesses comparable immunogenicity to KLH while owning a higher solubility after conjugation. SHC is extracted from the most popular seafood, the white-leg shrimp or *P. vannamei* in the Latin name. In our hand, from 1kg healthy shrimp, 50 mL hemolymph could be extracted and 1 g high-pure SHC could be acquired finally. This high yield could pave the way for SHC industrialization. In future, SHC would reduce the research and production costs related to vaccine and antibody development.

Except for *P. vannamei,* other crustaceans e.g. Argentine red shrimp (*Pleoticus muelleri*), Red swamp crawfish (*Procambarus clarkii*), Chinese mitten crab (*Eriocheir sinensis*), Giant tiger prawn (*Penaeus monodon*) etc. (15), also provide the resource for hemocyanin extraction. These crustaceans could be captured from wild areas or harvested by aquaculture. In this study, we selected the *P. vannamei,* given that this species dominates in the crustacean production worldwide. The high production of *P. vannamei* would guarantee the sustainable supply of shrimp hemocyanin at a relatively low expense. Furthermore, after hemolymph extraction, *P. vannamei* could be processed as a food product such as dried shrimp. Therefore, SHC is more economical than KLH.

In antibody development, SHC would be more suitable for hapten-conjugation than KLH considering the solubility. However, in vaccine research (19), especially for preparing human vaccines, one thing should be considered. *P. vannamei* is the most common seafood and is often put on the table, whether the anti-SHC antibody is present in the human body is not known. Pre-existing antibodies against SHC in people may reduce the vaccine potency or even induce allergy(20). Therefore, more studies are needed for applying SHC in vaccine research.

## Author contribution

Conceived and designed the project: B. Jiang.

Conducted the experiment: M. Wei extracted the hemolymph; H. Sun purified the SHC and performed the peptide conjugation assay; H. Sun & A. Guo performed the mouse-related assays; H. Sun & C, Zhang & R. Huang performed the WB detection assay; Y. Wang performed the IHC detection; A. Guo performed the hybridoma fusion and monoclonal antibody screening assays. XX. Li performed the size-exclusion chromatography and transmission electron-microscopy.

Drafted Manuscript and processed the data: B. Jiang & A. Guo.

Supervise the project: B. Jiang & J. Wu.

All authors discussed the manuscript.

## Acknowledgement

We thanks for Fuheng Biotech for their kindly gift of cell-lines and all other peoples who were involved in performing these experiments and gave us constructive advice.

## Patent information

The utilization of SHC in hapten-carrier preparation has been submitted to patent application. The patent application numbers in Republic of China were CN2023115993853, CN2024100341885. PCT patent number was PCT/CN2024/071375.

---

**Supplementary Figure 1.**
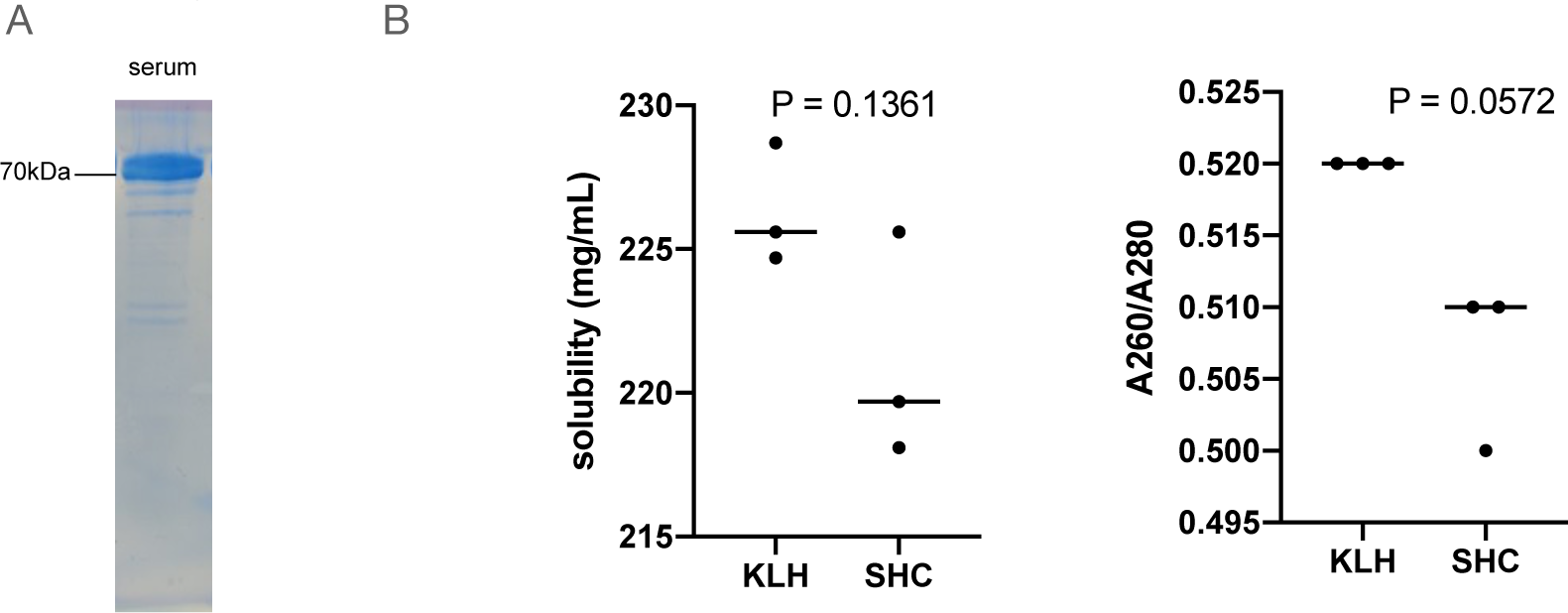

**Supplementary Figure 2.**
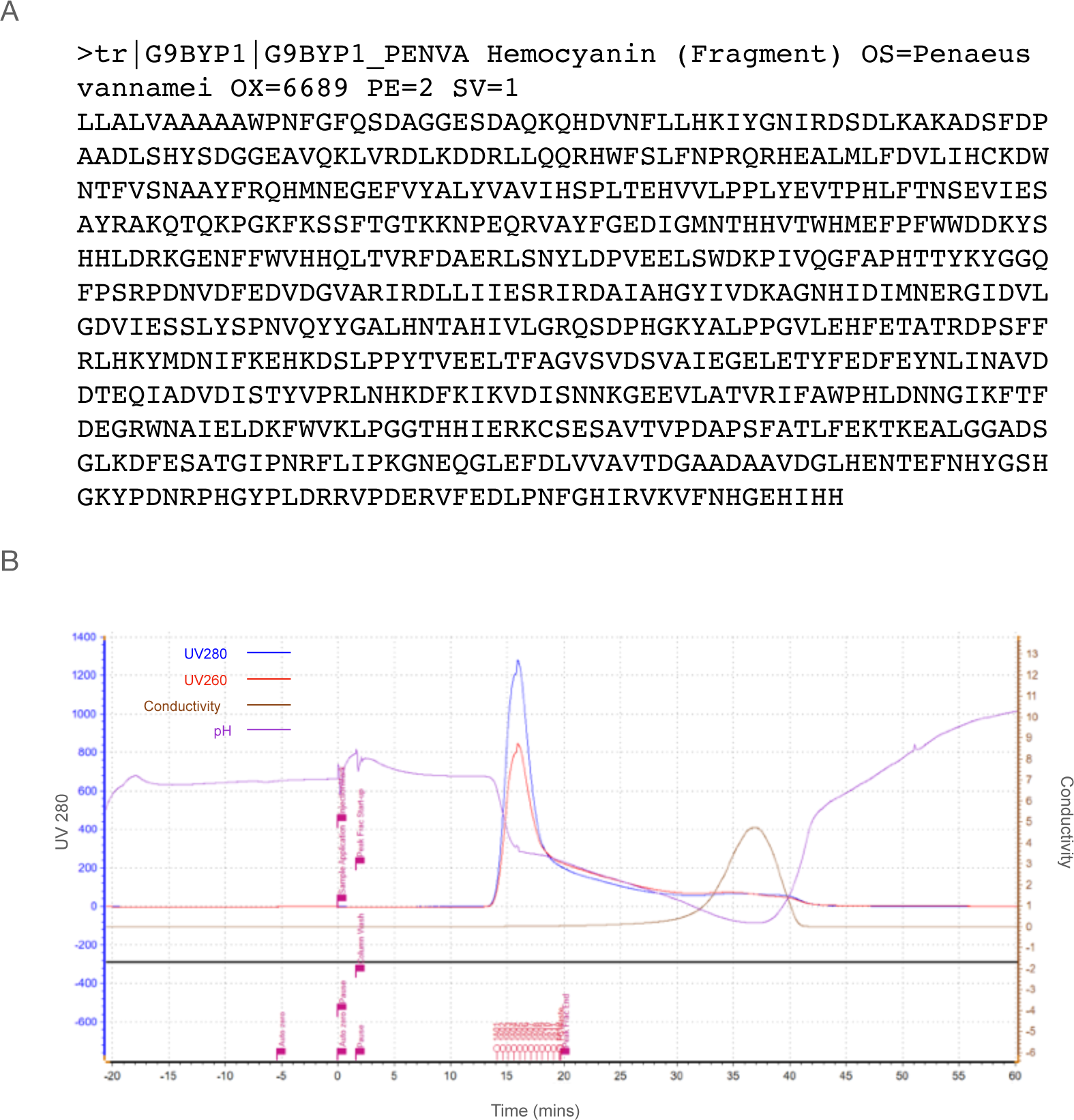

**Supplementary Figure 3.**
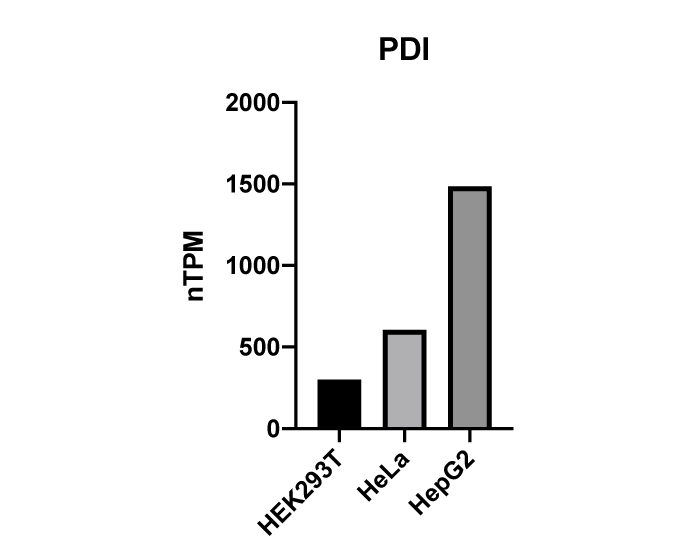

**Supplementary Figure 4.**
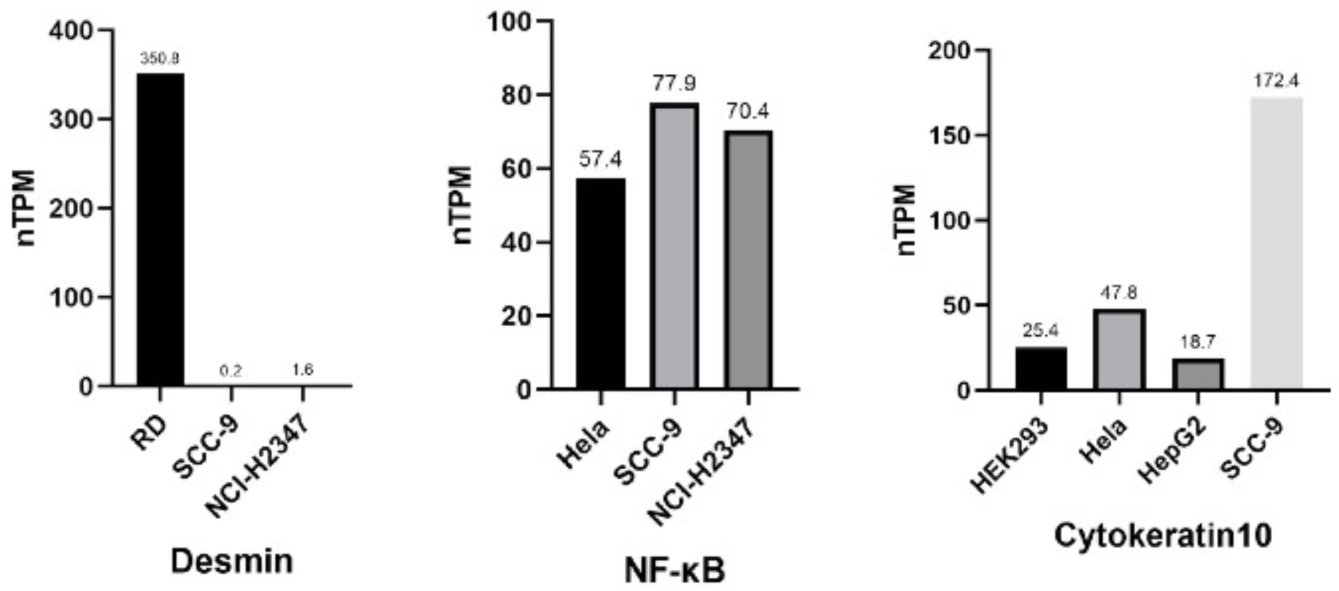

